# Where do the Dmanisi hominins fit on the human evolutionary tree?

**DOI:** 10.1101/2025.03.01.639363

**Authors:** Debbie Argue, José María Bermúdez de Castro, Michael S. Y. Lee, Maria Martinón-Torres

## Abstract

Archeological excavations at the site of Dmanisi in the Republic of Georgia have yielded a rich assemblage of hominin fossil remains, as well as lithic artefacts and bones of fossil fauna. The site is considered to be between 1.95 Ma and 1.77 Ma (Gabunia et al. 2000b) and presents us with the first skeletal evidence of hominins to emerge from Africa, a key event in human evolution. Their morphology, and the degree of morphological variation observed among the assemblage, has generated considerable controversy about their affinities and heterogeneity. Here we use parsimony analyses to test the competing hypotheses for Dmanisi hominins employing characters from the cranium, mandible, dentition, and postcranium; and we address anomalies in endocranial volume, dentition and mandibular structure among the assemblage. We propose that the Dmanisi hominins are not *Homo erectus*, and that two species are represented among the assemblage: one comprises *Homo georgicus* and the other an as yet unnamed species. Our review of the dating of the Dmanisi site leads us to propose that *Homo georgicus* was probably present by 1.8 Ma and that the other hominins recovered from the Dmanisi excavations accumulated at some time or times during the reverse polarity of 1.07 Ma and 1.77 Ma. The specific, individual, ages of these hominins remain unknown.

## 1. Introduction

The hominin remains recovered from the Dmanisi site comprise five crania, four of which have associated mandibles. Three of the mandibles include their teeth and postcranial remains have been recovered for two of the individuals. The assemblage was attributed to *Homo erectus* (Gabunia and Vekua 1995). When a second more complete and robust mandible, D2600 was discovered, a comparative analysis with *Homo habilis* and *Homo erectus* led Gabounia et al. (2002) to the recognition of a new *Homo* species: *H. georgicus*, with D2600 the holotype. Other taxonomic attributions have been proposed, however: *Homo erectus* (Rightmire et al. 2006); *H. sp. indet (aff ergaster)* (Rosas and Bermúdez de Castro 1998); *H. ex gr. ergaster* (Gabunia et al. 2000a); *Homo georgicus* (Gabounia et al. 2002); and *H. erectus ergaster georgicus* (Lordkipanidze et al. 2013). Vekua et al. (2012) deem the Dmanisi specimens to be the most primitive and small-brained fossils to be grouped with *H. erectus*, or any taxon linked unequivocally with genus *Homo,* and also the ones most similar to the presumed *habilis*-like stem.

The range of variation observed among the Dmanisi crania, mandibles and dentition has also prompted considerable discussion in the literature. Two potential explanations for the variability have been advanced: that the species exhibits an unusually large degree of sexual dimorphism; or that more than one species is contained among the sample.

Here we present parsimony analyses to assess the potential attributions of the Dmanisi hominins and assess whether the assemblage represents one or more species.

### 1.1. Background on fossil remains

The first hominin fossil of the Dmanisi assemblage to be discovered was a mandible, D211, recovered in 1991 in Block 1 of the excavations (Gabunia and Vekua 1995). The bone-bearing layer shows normal geomagnetic polarity indicating that the mandible dates to the end of the Olduvai Subchron (1.95-1.77 Myr) (Gabunia and Vekua, 1995).

In their comparative assessment of the mandible, Gabunia and Vekua (1995) observe that it is most similar to 1.7 million-year-old (mya) East African hominin KNM-ER 730; 1.5 mya KNM-ER 992 and KNM-WT 15000; ∼700,000 year old (Leakey and Row 1994:10); OH 22 from Olduvai Gorge, Tanzani;: and Middle Pleistocene Zhoukoudian G1-6 (China); and that it also shows similarities to members of a Tighenif/Arago/Mauer group, and a Sangiran group. Gabunia and Vekua (1995) propose that the most reasonable interpretation is that the mandible belonged to a population of *H. erectus*, possibly foreshadowing *H. sapiens*.

Motivated by the heterogeneous affinities of the mandible exemplified in Gabunia and Vekua’s (1995) assessment, Bräuer and Schultz (1996) undertake multivariate analyses to clarify its possible affinities. In some of these analyses, D211 clusters with early *Homo* (*H. habilis* and *H. rudolfensis*), but the authors also detect a number of derived traits. For example, they concluded that the symphyseal cross-section and small dental dimensions were more likely to occur in later *H. erectus* specimens than in early *Homo*. They also identify a number of characteristics, such as the anterior marginal tubercle and the rather well-developed chin region, that indicate D211 exhibits derived traits that are rarely found in early *H. erectu*s, and conclude that Dmanisi could represent a very progressive early *H. erectus*-like form with closer affinities to later *H. erectus*, or even archaic *H. sapiens*, than to other contemporaneous and younger *H. erectus* specimens from Africa and Indonesia (Bräuer and Schultz 1996).

Rosas and Bermúdez de Castro (1998) compared D211 with mandibles and teeth of *Australopithecus*, *H. habilis*, *H. ergaster, H. erectus* (Sangiran and Zhoukoudian), Atapuerca, Tighenif. *H. heidelbergensis*, *H. neanderthalensis*, and *H. sapiens* (Rosas and Bermúdez de Castro, 1998: Table 1), and proposed that the Dmanisi mandible exhibits a unique combination of primitive and derived traits. For example, the location of the anterior marginal tubercles in D211 approaches that found in early representatives of *Homo*; and the anterior location of the lateral prominence and the shape of the retromolar area in Dmanisi correspond to the primitive pattern of the corpus/ramus junction, approaching that found in early *Homo* specimens. Rosas and Bermúdez de Castro (1998) therefore considered the structure of the Dmanisi mandible as very primitive. However, they found that aspects of the dentition, such as the reduction of the talonid in the fourth premolars and a distally decreasing molar series seem to be derived, and are observed in specimens of *Homo ergaster* but differ from those generally present in later hominids. On the other hand, the morphology and the relatively small breadth of the Dmanisi anterior teeth and of P3, they suggest, show affinities of this specimen with early representatives of *Homo*, assigned either to *H. habilis* or to *H. ergaster*. The relatively small bucco-lingual dimensions of the lower incisors in the Zhoukoudian sample also suggest certain affinities between Asian *H. erectus* and Dmanisi.

**Table 1.** Specimens included in the analyses.

| Species and Institution | Skeletal component | Specimen |
| --- | --- | --- |
| ***Pan troglodytes***<br>Australian National University, Canberra; Australian Museum, Sydney; National Museum of Australia, Canberra (NMA items temporarily at ANU). | Cranium; cranial part | M6057 (male); M27547 (female); M32977 (female); M37431 (female); 2865 (NMA). |
| **Gorilla**<br>Australian National University, Canberra; Australian Museum, Sydney; National Museum of Australia, Canberra | Cranium; cranial part | M37431 (female); M43538 (male); M36149 (male); M36148 (male); Gorilla 4 (ANU); Gorilla 2864 (NMA); Gorilla 18789 (female; NMA) |
| (NMA items temporarily at ANU). |  |  |
| ***Pongo pygmaeus***<br>Australian National University, Canberra;<br>Australian Museum, Sydney; National Museum of Australia, Canberra<br>(NMA items temporarily at ANU). | Cranium; cranial part | M43539 (male); M3480 (female); M39946 (mature male but no crest); M39947 (male); M8608 (young adult male); S1639 (male); 2867 (NMA). |
| ***Homo floresiensis***<br>Indonesian National Centre for Archaeology (ARKENAS) | Cranium; cranial part;<br>postcranium<br>Mandible; lower dentition | LB1.<br>LB1; LB6. |
| ***Homo erectus***<br>Sangiran Museum and Centre for Geological Survey, Indonesia;<br>Senckenberg Museum, Frankfurt; Naturalis, Leiden. | Cranium; cranial part | Grogan Wetan; Trinil 1; Sangiran 2,3,4,12,17. |
|  | Mandible/lower dentition | Sangiran 1b, 9; 22; SB 8013; BK 7095; Hanoman 13. |
|  | Postcranium | Femur II, III, IV, V; Kresna II femur. |
| ***Homo habilis***<br>Kenya National Museums, Nairobi, Kenya; National Museum and House of Culture, Dar es Salaam, Tanzania. | Cranium; cranial part | KNM-ER 1813; OH 13; OH 16; OH 24. |
|  | Mandible/lower dentition | OH 7; KNM-ER 1501; KNM-ER 1502; KNM-ER 1507. |
|  | Postcranium | OH, 62; OH 20; OH 34; KNM-ER 1472; KNM-ER 1481. |
| ***Homo ergaster***<br>Kenya National Museums, Nairobi, Kenya | Cranium; cranial part | KNM-ER 730; KNM-ER 1808; KNM-WT 15000; KNM-ER 3733; KNM-ER 3883. |
|  | Mandible/lower dentition | KNM-ER 730; KNM-ER 992; KNM-ER 1808; KNM-WT 15000. |
|  | Postcranium | KNM-WT 15000. |
| ***Australopithecus afarensis***<br>National Museum of Ethiopia, Addis Ababa, Ethiopia | Cranium; cranial part | AL 58-22; AL 162-28; AL198-1; AL 199-1; AL 200-1; AL 333-1; AL 333-2; AL 333-105; AL 438-2; LH 29; BEL VP 1/1; EP 2400/00; Garusi. |
|  | Mandible/lower dentition | AL 128-23; AL 145-35; AL 188-1; AL 207-13; AL 266-1; AL 277-1; AL 333-43; AL 333-45; AL 74; AL 333-w-1; AL 333-w-12; AL 333-w-60; AL400-1A; AL 438-1; LH 2; LH 4. |
|  | Postcranium | AL 128-1; AL 129-1; AL 288-1 (cast); LH 21. |
| ***Australopithecus africanus***<br>University of Witwatersrand; Ditsong Museum. South Africa. | Cranium; cranial part | STS 5; STS 17; STS 52; STS 53; STW 13; STW 73; STW 151; STW 183; STW 505; TM 1511; TM 1512. |
|  | Mandible/lower dentition | STS 7; STS 36; STS 52; STW 151; STW 327; STW 384; STW 404; STW 498. |
|  | Postcranium | STS 14; STW 431; STW 992/100/101. |
| ***Australopithecus sediba***<br>University of Witwatersrand, Johannesburg, South Africa | Cranium; cranial part | MH 1. |
|  | Mandible/lower dentition | MH 1; MH 2. |
|  | Postcranium | MH 1; MH 2.<br>Elen Feuerriegel, pers. com. |
| ***Homo sapiens***<br>Australian National University, Canberra; Australian Museum, Sydney. | Crania; mandibles/dentition/postcrania | 37 male and female specimens from Africa (6, labelled ‘Negroid’); China (1); Europe (8); Indonesia (incl Liang Toge) (2); Jordan (3); Japan (10, incl 2 Ainu), ‘Melanesian’ (1), |
|  |  | New Ireland (1); Papua (2); Tahiti (1); Vietnam (1); Qafzeh skull II (cast). |
| **Dmanisi hominins**<br>Data obtained from the literature. | Crania, mandibles; dentition; postcrania | **Cranium and associated mandible:**<br>**D2282 + D211**<br>Bermúdez de Castro et al. (2014)<br>Bräuer and Schultz (1996)<br>Gabunia and Vekua (1995)<br>Martin-Francés et al. (2014)<br>Rightmire et al. (2006)<br>Rosas and Bermúdez de Castro (1998)<br>Van Arsdale (2006)<br>**D2700 + D2735**<br>Bermúdez de Castro et al. (2014)<br>Martin-Francés et al. (2014)<br>Rightmire et al. (2006)<br>Van Arsdale (2006)<br>**D4500 + D2600**<br>Bermúdez de Castro et al. (2014)<br>Lordkipanidze et al. (2013)<br>Van Arsdale (2006)<br>**Cranium D2280**<br>Gabunia et al. (2000b)<br>Rightmire et al. (2006)<br>**Dentition:**<br>Martinón-Torres et al. (2008)<br>Rightmire et al. (2006)<br>Rosas and Bermúdez de Castro (1998)<br>Van Arsdale (2006)<br>**Postcranium:**<br>Lordkipanidze et al. (2007)<br>Pontzer et al. (2010) |
|  |  | Elen Feuerriegel, character states for characters 81, 82, 83, 84, 104 (pers. comm.). |
| *Homo naledi* | Cranium, mandible, postcranium, dentition | Berger et al. (2015)<br>Elen Feuerriegel (pers. comm.)<br>Hawks et al. (2017)<br>Laird et al. (2017). |

Rosas and Bermúdez de Castro (1998) propose that the Dmanisi mandible might be taxonomically classified as *Homo sp. indet.* (*aff. ergaster*).

Two partial crania, D2280 and D2282, were recovered in 1999 at ‘almost the same stratigraphic level’ (Gabunia et al. 2000a; p.1022) in which mandible D211 had earlier been discovered (Gabunia and Vekua 1995). D2282 represents the same individual as mandible D211. Gabunia et al. (2000a) assign D2280 and D2282_ D211 to *Homo ex gr. ergaster*.

Schwartz (2000), however, views the differences between the crania as reflecting different species rather than intrapopulation differences. They cite differences in the supraorbital, mastoid and nuchal areas, in the course of the vault sutures; in cranial outline and cross section; and cranial size.

The discovery of a third cranium, D2700, recovered from Block 2 of the excavations, located about 10m west of Block 1 (Ferring et al. 2011; Figure 1) generated further questions about heterogeneity in the assemblage. Lee (2005) tested whether the variation in cranial capacities in the fossil sample, D2280 (775 cc; Gabunia et al. 2000a); D2282 (estimated to be 650 cc; Gabunia et al. 2000a); and D2700 (600 cc; Vekua et al. 2002), exceeds that within modern humans, chimpanzees, and gorillas. Treating the larger D2280 as male and the other two as females, as hypothesised by Gabunia et al. (2000a), Lee (2005) found that variation observed in the Dmanisi sample can easily be found in male-female pairs of modern humans, chimpanzees, and gorillas; and concludes that the null hypothesis of a single species cannot be rejected. However, if all crania represent females, Lee considers that it is unlikely that a pair of females could generate the level of difference in the Dmanisi sample (Lee 2005).

**Fig 1.**
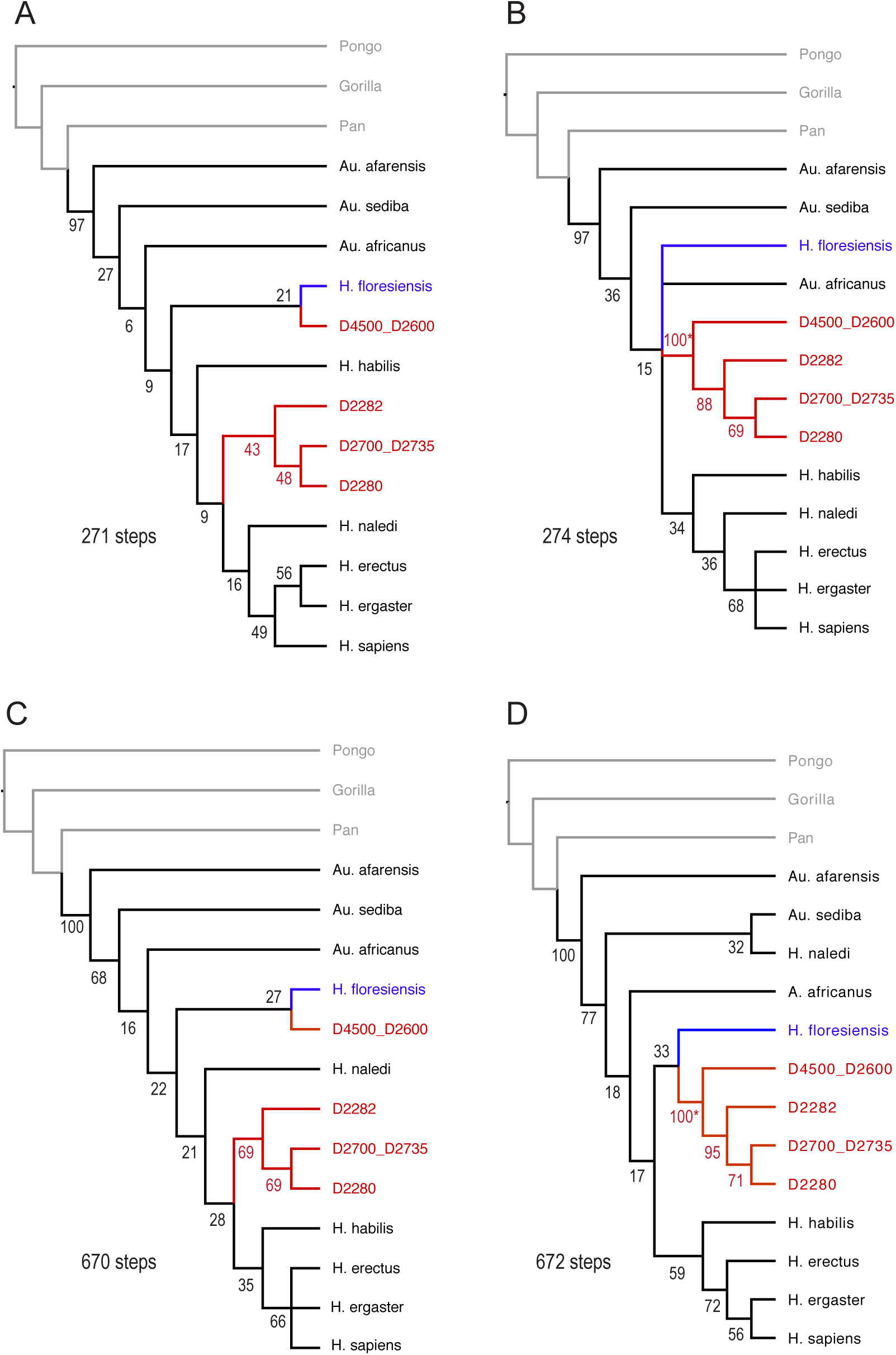
Phylogenetic hypotheses for Dmanisi hominins, based on parsimony analysis employing (A) multistate coding, without constraints; (B) multistate coding, with Dmanisi individuals constrained to form a clade; (C) intermediate coding, without constraints; (D) intermediate coding, with Dmanisi individuals constrained to form a clade. Dmanisi individuals in red, non-hominin outgroups in grey; numbers refer to bootstrap support.

In 2000 a second mandible, D2600, discovered in volcanic ashes in Block 2 of the excavations, is dated to 1.81 +/-0.08 Ma (de Lumley et al. 2002; Gabounia et al. 2002), although this date has been disputed by Ferring et al. (2022), (see below). The dimensions of D2600 are far greater than mandible D211. From a comparative analysis of D2600, D211 and mandibles of *Au. africanus*, *Au. boisei* and *Au. robustus (*now attributed to *Paranthropus*); *H. habilis*; Sangiran *H. erectus*; Zhoukoutian *H. erectus*; *H. ergaster* KNM-WT 15000; KNM-ER 992; and *H. neanderthalensis*, Gabunia et al. (2002) raise the possibility that two groups coexisted at Dmanisi. After considering this, however, and examining a range of characteristics on the two Dmanisi mandibles, they propose that the mandibles share a unique suite of shared characteristics. They performed a formal diagnosis, establishing a new species, *Homo georgicus*, sp. nov., that expresses a marked degree of sexual dimorphism (Gabounia et al. 2002).

Rightmire et al. (2006) attributed the variation among the Dmanisi crania and mandibles as probably related to growth stage and sexual dimorphism, and they consider it appropriate to group the Dmanisi hominins together as one paleodeme (species), that is close to the stem from which *H. erectus* evolved. They express some reservation, however, as to whether D2600 should be included, acknowledging that they have not reached a consensus: one author regards the large mandible D2600 as different from others, and prefers to include it in *H. georgicus*.

Van Arsdale (2006) tested the null hypothesis that the dimorphism observed among the Dmanisi mandibles D211, D2735 (associated with cranium D2700) and D2600 is the result of a process of sampling different age and different sex individuals from a single evolutionary group. He employs a random sampling strategy using 31 linear measures from the Dmanisi mandibles, *H. sapiens*, *Pan troglodytes* and *Gorilla gorilla gorilla*. The measurements comprise aspects of corpus height, breadth, tooth size, dental arcade size and shape, symphysis size and shape, and ramus breadth. The amount of variation among the Dmanisi hominins, he finds, exceeds that found among *H. sapiens* and *Pan* and he rejected the single-species hypothesis for the Dmanisi hominins. When compared with a *Gorilla* model, however, the observed Dmanisi variation does not exceed the expected level of variation, except for the height of the mandible at M3. Overall, Van Arsdale concludes that the results support the view that the variation within the Dmanisi sample is not consistent with a mixed-taxa sample on the basis of the comparative materials available. Rather, he proposes that the Dmanisi hominins comprise a single hominid taxon with greater than expected sexual dimorphism than is evident in modern humans or chimpanzees. Thus, the null hypothesis may not hold when humans or chimpanzees are used as reference populations, but it cannot be rejected for a model based on the more highly dimorphic gorilla (Van Arsdale 2006).

To assess size variation among the Dmanisi mandibles, Skinner et al. (2006) used shape analyses based on four linear variables. Their results lead them to propose that the degree of sexual dimorphism evident among the Dmanisi mandibles exceeds that of other species of *Homo*; and is rarely seen in any extant hominoid species. They present two hypotheses: i) the degree of sexual dimorphism in the Dmanisi taxon exceeds expectations, thus highlighting the need to reconsider conclusions about its inclusion in, and/or the definition of, the genus *Homo* (Skinner et al. 2006:45). Their other hypothesis is that the presence of a second hominin species, represented by D2600, should be considered, on the basis that D2600 is from a different stratigraphic layer than D211, possesses morphological features not seen in the other Dmanisi mandibles (e.g., different molar size gradient), and that the rest of the Dmanisi fossils do not provide strong evidence of a highly sexually dimorphic species (Skinner et al. 2006:45).

Partly in response to Skinner et al.,’s (2006) use of only four variables, Rightmire et al. (2008), employed a random sampling strategy on 26 measurements for intact and relatively undamaged portions of the D211 and D2600 mandibles and their dentitions. They found similar results to those of Van Arsdale (2006): the degree of variation between D211 and D2600 is outside the range for recent humans and, in some cases, also excessive compared to chimpanzees. When a gorilla model of variation is applied, only one of the Dmanisi ratios (not identified) suggested significantly more variability than expected. They conclude that the Dmanisi hominins differ no more than would be expected for individuals within a relatively dimorphic ape population and that there are currently no compelling anatomical grounds for sorting any of the Dmanisi fossils to other than a single species.

Variation in the dental remains in Dmanisi mandibles has also generated discussion. Martinón-Torres et al. (2008) found that D2600 retains a number of primitive conditions, including the primitive molar sequence M1 < M2 < M3, also observed in *Australopithecus*, *Paranthropus*, and the majority of *H. habilis*, *H. rudolfensis*, and the early *Homo* dental remains. In contrast, D211 and D2735, exhibit the first occurrence of M1 > M2 (Martinón-Torres et al. 2008). This is a characteristic that does not occur again until the Middle Pleistocene, when it occurs only infrequently, observed in specimens such as OH 22, Middle Pleistocene specimen from Africa, Rabat, as well as in some Atapuerca-Sima de los Huesos hominins (Spain) and Zhoukoudian individuals (Martinón-Torres et al. 2008). The occurrence of M1>M2 is, then, a peculiar trait for the geological time frame of the Dmanisi hominins (Martinón-Torres et al. 2008). The third molar is missing from D2735, but in D211 the M2/M3 sequence is M2 >M3, occurring more commonly than M1>M2, and is also found in OH 16, KNM-ER 806, KNM-ER 730, and KNM-ER 992, and some other specimens assigned to *H. erectus* and *H. antecessor*. In contrast, D2600 shares the primitive pattern (M2 < M3) with *Australopithecus*, *Paranthropus*, and the majority of the early *Homo* specimens (Martinón-Torres et al. 2008).

The lower third premolars of D2600 possess distinct mesial and distal roots that bifurcate close to the cemento-enamel junction as occurs in KNM-ER 1802 and the 2.5 – 2.3 mya Uraha mandible (UR-501) from Northern Malawi, Africa, as well as in *P. robustus* and *P. boisei* (Martinón-Torres et al. 2008). Martinón-Torres et al. (2008) observe that this type of molarized bifurcation is not typical of the *Homo* fossil record, where double-rooted P3s tend to be of the Tomes’ type. The differences between the premolar root morphology, molar sequences and dental dimensions on the two smaller Dmanisi mandibles compared to that on D2600 suggested to Martinón-Torres et al. (2008) that two paleodemes are present among the Dmanisi hominins. Schwartz et al. (2014) also discuss species-distinctive differences in the dentition in D2600 compared to that of the other Dmanisi hominins, citing aspects of the lower canines, premolars, and M3, P3 root form, all of which, they argue, exceed the differences found in *H. sapiens*.

Bermúdez de Castro et al. (2014) observed that the Dmanisi mandibles D211, D2735 and D2600 present a set of seven common primitive traits for the *Homo* clade: mental foramen placed at the level of P3–P4; presence of mental protuberance; anterior marginal tubercle at the level of the canine or the C/P3; U-shaped arcade: high value for the alveolar arcade index; shift between the anterior and posterior parts of the dental arcade at the level of the canine; conspicuous alveolar prominence; and a prominent superior transverse torus (Bermúdez de Castro et al. 2014: Table 4). D211 and D2735, however share features that are different from D2600, each of which is established early in the ontogeny (during childhood or even before) and that do not depend on size or sexual dimorphism (Bermúdez de Castro et al. 2014). Early in ontogeny, when dental development is still in progress, the differences in mandibular shape at the genus level and at the species level are already established. Mandibular shape differences are reached near or before birth; and the spatial relationships between component parts (e.g. ramus and corpus) are established early in ontogeny and remain invariant during growth (Bermúdez de Castro et al. 2014). Bermúdez de Castro et al. (2014) observe a distinctive and primitive spatial relationship between the corpus and the ramus on D2735 (and apparently in D211 as well), in which the plane of the ramus is positioned buccally in relation to the corpus. This morphology is found on mandibles of *Au. anamensis*, *Au. afarensis*, *Au. africanus*, *P. africanus*, *P. boisei* and *H. habilis*, but is not present in D2600 (Bermúdez de Castro et al. 2014). Other differential features between D2600 and the two smaller Dmanisi mandibles include the presence of lateral tubercles near the mental protuberance in D2600 and its exaggerated height of the corpus at this level (49.0 mm), in contrast to 30.8 mm in D211 and 32.3 mm in D2735. The marked projection of the inferior transverse torus in D2600 is visible in superior view, whereas the inferior transverse torus in D211 and D2735 is hidden by the superior transverse torus in superior view. The myohyloid line is diffuse in D211 and D2735, but conspicuous on D2600, and the extramolar sulcus is relatively wide in D211 and D2735 but narrow on D2600. In all, Bermúdez de Castro et al. (2014) observe 10 features on D2600 that are not shared with the other two Dmanisi individuals. Considering the suite of differences between D2600 the other two Dmanisi mandibles, Bermúdez de Castro et al. (2014) suggest that two paleodemes (species) are present among the Dmanisi hominins: *H. georgicus*, represented by D2600; and a separate paleodeme (species) represented by D211 and D2735 that has a completely different cranio-facial growth pattern to D4500_D2600.

Baab (2008), examining the taxonomic status of *H. erectus* sensu lato using 3D geometric morphometric data, and a broad range of human and nonhuman primate samples, test whether variation in *H. erectus* s. l. most clearly resembles that seen in one or more species. The Dmanisi cranium D2280 is included in the comparative sample. Baab’s (2008) first analysis focuses on 16 landmarks that are available for the reference taxa: humans, apes, and both extant and extinct papionin monkeys, and crania ascribed to *H. erectus* s. l. in this analysis: KNM-ER 3733, KNM-ER 3883, Olduvai Gorge OH 9, Bouri, Daka, Sangiran 17, four Ngandong crania, Sambungmacan 3; and four Zhoukoudan crania. In this analysis, Dmanisi D2280 is closest to KNM-ER 3883 in cranial vault shape. When performing a second analysis in which a 32-landmark set designed to encompasses a greater range of cranial landmarks, that in this case is available only for the fossil and extant hominins, Baab (2008) finds that D2280 is most similar in cranial vault shape to KNM-ER 3733 and KNM-ER 3883 (Baab 2008). Baab (2008) interprets the results to most strongly support a single-species interpretation for *H. erectus* s. l. that includes the Dmanisi individual represented by D2280.

Another cranium, D4500, recovered in 2005 from Block 2 in the Dmanisi excavations, represents the same individual as mandible D2600 (Lordkipanidze et al. 2013). D4500’s endocranial capacity is 546 +/-5 cm^3^ (Lordkipanidze et al. 2013), notably smaller than the other Dmanisi crania and only marginally larger than Stw 505 (*Au. africanus*) from Sterkfontein, and essentially the same as AL 444-2 (*Au. afarensis)* from Hadar (Rightmire et al. 2017). D4500’s skull is low, but comparatively wide and elongate, with a frontal that slopes moderately behind a thick, bar-like supraorbital torus. The glabellar region is massive and substantially overhangs the mid-face. Postorbital constriction is marked. The face is flanked by massive zygomatic arches, and spacious temporal fossae indicate large masticatory muscles. The palate is relatively narrow and the mandibular symphysis deep (Rightmire et al. 2017). Its muzzle-like face is similar to *Au. afarensis* and is among the largest and most prognathic known from early *Homo* (Lordkipanidze et al. 2013). Overall, Lordkipanidze et al. (2013) view the cranial shape variation within early *Homo* paleodemes as similar in mode and range to that seen within modern *Pan* demes and that ‘the morphological diversity in the African fossil *Homo* record around 1.8 Ma probably reflects variation between demes of a single evolving lineage, which is appropriately named *H. erectus*’ (Lordkipanidze et al. 2013; p. 330).

Baab (2016) performs a 3D analysis of hominins *H. erectus* s. l. In the analysis D2289, D2282, D3444, D2280 and D2700 generally fell in or near the range of variation of the *H. erectus* s. l. sample, and were more similar in overall vault shape to the Koobi Fora crania KNM-ER 3733 and KNM-ER 3883. Baab (2016) proposes that the Dmanisi hominins should probably be included in *H. erectus* s. l., observing that the Dmanisi fossils expand the variation in that species. Baab (2016) comments, however, that D3444, recovered from Block 2, and the newly discovered cranium Dmanisi D4500 (not included in the analyses) are the most robust of the five Dmanisi crania and yet are quite different in shape from one another, and that it is unlikely that either pattern can be entirely attributed to sexual variation, although this issue is not discussed further.

Recognising that the relationship of D4500 to other Dmanisi individuals is a key issue, not yet resolved, Rightmire et al. (2017) evaluate the null hypothesis that only one taxon is represented at the site. Using the coefficient of variation (CV) to compare Dmanisi with appropriate reference groups including recent humans and fossil hominins. They examine the characteristics of the Dmanisi crania in relation to *Au. afarensis*, *Au. africanus*, *Au. sediba*, *H. habilis*, *H. rudolfensis, H. naledi* and *H. erectus*. They identify the presence of *H. habilis* traits, as well as traits diagnostic for *H. erectus* in the Dmanisi crania, noting that it is difficult to find diagnostic traits by which the Dmanisi hominins can be sorted. They conclude that the Dmanisi hominins cannot be attributed to either *H. habilis* or *H. erectus*. Rather, they propose that *H. habilis*, the Dmanisi group, and *H. erectus* constitute segments of a single evolutionary lineage (Rightmire et al. 2017).

In summary, there are conflicting views about the attribution of the Dmanisi hominins, and the variation among them has led to questions as to their heterogeneity. Here we use parsimony analyses to test the competing hypotheses for Dmanisi hominins employing characters from the cranium, mandible, dentition, and postcranium; and we address cranial, mandibular and dental anomalies among the assemblage.

### 1.2. Dating and stratigraphy

The chronology of the site of Dmanisi is of paramount importance for our understanding of hominin movement out of Africa. Excavations at Dmanisi have revealed a sequence of strata above the 80m thick Mashavera Basalt lava flow. The basalt has been dated to 1.8 ± 0.1 Ma by K/Ar dating, 2.0 ± 0.01 by ^40^Ar/^39^ (Gabunia et al. 2000a) and Ferring et al. (2022) report dates from 4 A/Ar samples from the Mashavera Basalt that range between 1.77 ± 0.04 and 1.79 ± 0.04 (Ferring et al. 2022; SOM Table 1). Because the ^40^Ar/^39^Ar ages and the K/Ar ages are equivalent at the 2s level confidence, Ferring et al. (2022) pool all 6 ages and calculate an age of 1.799 ± 0.014 Ma for the eruption which delivered the Mashavera Basalt.

The strata overlying the Mashavera Basalt comprise Stratum A, immediately above the Mashavera Basalt, and Strata B1-B5 (Ferring et al. 2022).

The hominin fossils are contained within a complex series of pipes, comprising eroded subsurface tunnels and cavities that incise Stratum A. At some time or times, fluvial action transported hominin and faunal bones from the overlying Stratum B1 into the pipes, where the remains, judging by their condition, were quickly encased with sediment (Vekua et al. 2002; Lordkipanidze et al. 2013; Rightmire et al. 2017). Stratum B1 and the pipes are sealed off by Stratum B2. Above Stratum B2 lie strata B3 to B5 (Ferring et al. 2022). We do not know the absolute ages of these strata, nor their various rates of formation; nor do we know when pipe onset or infill events occurred.

Stratum B is reversed polarity, correlated with the upper (younger) part of the Matuyama Chron (Coil et al. 2020; Ferring et al. 2022). We infer from Ferring et al. (2022) that Stratum B is younger than the Olduvai Subchron which is dated to 1.78 Ma-1.95 Ma (Lourens et al. 1996) and older than Jaramillo Subchron (0.95 Ma-1.05 Ma). Stratum B is therefore within the range of 1.05 Ma to 1.78 Ma. The inference, then, is that hominin remains recovered from the geological pipes that emanate from Stratum B1 could have been deposited in Stratum B1 anytime, or times, between 1.05 and 1.78 Ma. Agusti et al. (2022), however, have identified a tooth of *Mimomys pliocaenicus* among the small mammals recovered from the base of Stratum B1. *M. pliocaenicus* is a small vole that is a characteristic element of the Late Pliocene (late Villanyian; MN17) faunas from Europe (Agusti et al. 2022). The Late Villanyian is dated to between 2.58 – 1.806 million years ago (Uhen et al. 2023). The presence of this species among the faunal remains implies a basal date for Stratum B of 1.8 Ma, which seems at odds with the reverse polarity determination for Stratum B.

Soon after the discovery of the first mandible, D211, tentatively inferred to be 1.95 – 1.77 Ma (Gabounia and Vekua 1995), Braüer and Shultz (1996) observed that a typical representative of *Mammuthus* (*Archidiskodon*) *meridionalis* is present in the faunal assemblage from the site excavations. That species, they note, appears as late as the earliest Middle Pleistocene and that ‘the existence of a typical representative of *A. meridionalis* in Dmanisi does not necessarily indicate an age of more than 1.5 Ma (Ubeidiya) because this elephant species still appears as late as the earliest Middle Pleistocene’ (Bräuer and Schultz 1996; p. 446). That is, Bräuer and Schultz (1996) suggest that the D211 mandible might be considerably younger than 1.95 Ma-1.77 Ma inferred by Gabunia and Vekua (1995).

Four years later, Goguitchaichvili and Pares (2000) performed palaeomagnetic and rock-magnetic studies, revealing the presence of reverse magnetisations in the sediments from which mandible D211 and crania D2280 and D2282 had been recovered. The reverse magnetisation of the sediment led Goguitchaichvili and Pares (2000) to propose that the Dmanisi site most likely has an age between 1.07 Ma and 1.77 Ma.

Based on the paleomagnetic and geochronological data, Gabunia et al. (2000a) also conclude that the fossil-bearing sedimentary rocks at Dmanisi are reverse polarity and are thus constrained within the Matuyama chron, calibrated between 1.07 Ma and 1.77 Ma. They argue, however, that the Oldowan, or Mode 1 lithic assemblage, and, in particular, the age of the majority of the associated vertebrate fauna, indicate a latest Pliocene, Olduvai Subchron age of 1.95 to 1.77 Ma. They conclude that this evidence restricts the age of the remains to slightly younger than the Olduvai-Matuyama boundary, or about 1.77 Ma (Gabunia et al. 2000a). 1.77 Ma has since become the most often cited date for the Dmanisi fossil hominins.

Some of faunal species recovered from the Dmanisi excavations, however, suggest a different age determination for the hominin fossils. Although the majority of the associated fossil fauna in the Dmanisi assemblage suggests that the hominins date to around 1.77 Ma, Lordkipanidze et al. (2007; S2) observe that more modern forms than those from Late Pliocene contexts are represented, listing *Cervus* (*Pseudodama*) cf. *nestii*; *Bison* (*Eobison*) *georgicus*; and *Pontoceros* sp. The latter is now attributed to a new species, *Pontoceros surpine* sp. nov. Vekua, 2012-03-12 (Vekua 2012). Gabunia et al.’s (2000b) compilation of fauna from the Dmanisi excavations contains *Soergelia* (Gabunia et al. 2000b; p. 1024) and Tappen et al. (2022) include hyena *Pachycrocuta brevirostris* and *Stephanorhinus* in their faunal list (Tappen et al. 2022; Suppl Table S1).

*Pontoceros, Bison, Soergelia, Pachycrocuta brevirostris* and *Stephanorhinus* are among a suite of fauna that dispersed to Europe between 2.0 and 1.8 Ma where they displaced the typical Villafranchian faunas of the Eastern Mediterranean and Europe (Koufos et al. 2005). They are known until at least a million years ago, with *Soergelia* continuing to less than a million years ago (Koufos et al. 2005), while *Pachycrocuta brevirostris* existed until at least 0.78 Ma; and *Archidiskodon meridianalis’*s last known appearance is between 0.9 and 0.78 Ma (Vislobokova and Agadjanian 2015; Table 2). *Bison* (*Eobison*) appeared 1.8 Ma, remaining until between 1.2 and 1.1 Ma (Vislobokova and Agadjanian 2015; Table 2). *Stephanorhinus*, which is represented by an entire skull that was associated with hominin skull D2700 and its mandible D2735, has been identified as *Stephanorhinus etruscus etruscus* (Vekua et al. 2002). Among the Eurasian rhinoceroses, *Stephanorhinus etruscus* is one of the most recorded and widespread extinct species of the genus. *S. etruscus* existed from 3.3 Ma to ca. 0.90 Ma (Pandolfi et al. 2017). There are then at least eight species represented in the Dmanisi assemblage that persisted beyond the Latest Pliocene. This fauna could have frequented the Dmanisi region at any time, or times, during the period of their existence, their remains accumulating in the Dmanisi strata at unknown periods, although within the reverse polarity dates for the pipe sediments, that is, between 1.78 and 1.07 Ma. That is, the Dmanisi hominin fossils, with which the fauna was associated (Tappen et al. 2022), could have potentially accumulated at any time, or at various times, during the 700,000-year period of reverse polarity. The veracity of a single point age estimate of 1.77 Ma for the site and the hominins, then, requires further elucidation.

**Table 2.** Attribution of Dmanisi hominin postcrania to individuals, following Lordkipanidze et al. (2007; p. 306).

| Cranium | Associated mandible | Associated postcrania |
| --- | --- | --- |
| D2282 | D211 | - |
| D2700 | D2735 | D2724 left clavicle<br>D2715 right humerus<br>D2680 left humerus<br>D3160 left femur fragment |
| D4500 | D2600 (*H. georgicus* holotype) | D4162/D4161 right/left clavicles<br>D4507 left humerus<br>D4167 right femur<br>D3901 right tibia |
| D2280 | - | - |

Mandible D2600 was found in volcanic ash in stratum B1y, the lowest of Stratum B1, immediately overlying the Mashavera Basalt. De Lumley et al. (2002) performed the first direct dating of the ashes in which the mandible was discovered. Using the dating method ^40^Ar/^39^Ar on volcanic glass fragments contained in the ash yielded an age of 1.81 ± 0.05 Ma (de Lumley et al. 2002). Subsequently, Garcia et al. (2010) sampled ash from two locations at the Dmanisi site. One sample was taken directly beside D2600 (Site A; Garcia et al. 2010; Figure 3; p. 445); and ten other samples were taken from Site B, a pit dug about 15 m northwest Site A (Garcia et al. 2010; and see Figures 3, 4). Based on ^40^Ar/^39^Ar data obtained by analysing glass fractions from these samples, Garcia et al. (2010) obtain an age of 1.81 ± 0.03 Ma for the ash. As the ash appears to be contemporary with the human remains, they propose that this age represents the age of the hominin represented by D2600 and its associated bones. As well, these ashes display normal polarity correlated with the Olduvai Subchron (Gabunia et al. 2000a; Calvo-Rathert et al. 2008; Garcia et al. 2010; Ferring et al. 2022). Ferring et al. (2022) reject de Lumley et al.’s (2002) and Garcia et al.’s (2010) dating based upon several potential methodological challenges when applying ^40^Ar/^39^Ar analyses to glass fragments (Ferring et al. 2022; SOM S1).

Above Stratum B1y lies Stratum B1x. This stratum contains subadult cranium D2700, its mandible, D2735, and postcranial elements. The remains are stratigraphically separated from the stratum below (B1y) (Lordkipanidze et al. 2007; and see Ferring et al. 2022 and Ferring et al. 2022; SOM). An adult individual represented by a single metatarsal was found at the higher stratigraphic position, Stratum B1z (Lordipanidze et al. 2007; Bermúdez de Castro et al. 2014). We do not have dates for strata B1x or B1z, so we do not know the absolute date for the individual D2700_D2735.

It is not yet clear how the stratigraphy in Block 1 relates to that of Block 2 strata. Whether Block 1 is part of the hominin-bearing pipe in Block 2, or if the Block 1 pipe represents different erosional events, does not appear to have been discussed in the literature. There are no dates available yet for the Block 1 excavations, so we do not know the dates for D211, D2280 or D2282.

Ferring et al. (2011), excavating in the M5 unit of Dmanisi 100 m to the west of the main excavations, recovered 73 artifacts from stratum A over a vertical range of ∼1.5 m. No hominin remains were found. The artifacts first appear in the lowest part of stratum A2, in level A2a, and provide evidence of the first occupations of the area. A2a stratum lies immediately above ∼30cm layer of ash. Below this ash is the Mashavera Basalt. The age of the Mashavera Basalt is used to infer the age of the artefacts. At the time of the discovery of the artefacts, the magnetostratigraphy and ^40^Ar/^39^Ar dating of the Mashavera Basalt was between 1.85 Ma and 1.78 Ma (Ferring et al. 2011). Ferring et al. (2011) infer that the earliest presence of hominins at Dmanisi occurred shortly after 1.85 Ma, and continued over the last half of the Olduvai subchron, 1.85–1.78. Ma. The Mashavera Basalt, however, has since been re-dated to 1.799 ± 0.014 Ma (Ferring et al. 2022). It is possible, then, that the earliest known presence of hominins at Dmanisi was after 1.799 Ma. A more honed date for the earliest presence at Dmanisi, represented by the lithic artifacts, may possibly be obtained if the 30cm band of A1 ashes that form a stratum between the Mashavera Basalt and the artefact-bearing strata were to be dated, rather than relying on inferences from dating the basal Mashavera Basalt.

The dating of the Dmanisi strata appears ambiguous and requires re-examination. There are horizontally extensive ashfalls in Stratum B1 (Lordkipanidze et al. 2007; Supplementary S1, p. 1); and in Strata B2, B3 and B4 (Coil et al. 2020). Rapid burial of hominin remains by serial episodes of ash fall deposition and the subterranean piping in Block 1 and Block 2, led to the near perfect preservation of the hominin remains (Lordkipanidze et al. 2006; Rightmire et al. 2017). The sediments from which D2700_D2735 and D2282_D211 were excavated are largely comprised of volcanic ash (Pontzer et al. 2011).

Dating the ashes that occur with the fossil hominins in the pipe formations could illuminate their age. Dating the ashfalls in Strata B1, from whence the fossil hominins and faunal remains came, would provide an approximation of the lower chronological limit for the period of fossil accumulation. Dating the ash in Stratum B2, that sealed in the B1 deposits, would provide an upper chronological limit.

## 2. Materials and Methods

### 2.1 Materials

This research is based upon the anatomical comparison of cranial, dental and postcranial features of the Dmanisi hominins with those of *Australopithecus* and *Homo*. We perform a cladistic analysis in which our comparative sample (Table 1) comprises the Dmanisi hominins, *Australopithecus afarensis*, *Au. africanus*, *Au. sediba*; *Homo habilis*; *H. erectus*; *H. ergaster*; *H. naledi*; *H. floresiensis*; and *H. sapiens*. *Pan, Gorilla* and *Pongo* are included as outgroups to identify ancestral, or pleisiomorphic, states for *Australopithecus* and *Homo*.

To test the affinities of the Dmanisi hominins, and whether they comprise one or more species, we treat each individual represented in the assemblage as a separate terminal taxon (Table 2). The Dmanisi individual comprising cranium D3444 and mandible D3900 exhibits extensive bone loss on the maxillae and the mandible due to resorption. Substantial tooth loss probably caused the bone remodelling several years before death (Lordkipanidze et al. 2006). Some of the character states for D3444 and D3900, therefore, are likely to be compromised, and we exclude this individual from our analyses.

Our characters (Supplementary Online Material SOM S1; SOM S2) and data are based on Argue et al. (2017) who obtained data from the original bones and fossils of *Au. afarensis*, *Au. africanus*, *Au. sediba*, *H. floresiensis*, *H. erectus*, *H. habilis*, *H. sapiens*, *Pan*, *Gorilla* and *Pongo* supplemented by data from the literature (Table 1).

We calculated indices designed to express aspects of cranial shape: porion-vertex height in relation to cranial breadth; relative biporionic width in relation to bi-parietal breadth; and internal length of palate in relation to width. Metric data sources for characters based upon indices (characters 1, 50, 52) come from Berger et al. (2015), Bermúdez de Castro et al. (2014), Bermúdez de Castro (pers. comm.), Braga et al. (1998), Brown et al. (2004), Brown and Maeda (2009), de Ruiter et al. (2019), Falk (1993), Groves and Argue (2010), Irish et al. (2013), Kaifu et al. (2005), Kimbel et al. (2004), Laird et al. (2017), Lordkipanidze et al. (2013), Rightmire et al. (2017), Tobias (1991), Walker and Leakey (1993), Wood et al. (1988), and Wood (1991). Other character state data sources are listed in Table 1.

We allocate the Dmanisi postcranial data to the two individuals D4500_D2600 and D2700_D2735, as designated by Lordkipanidze et al. (2007; p. 306). Our postcranial data are from Lordkipanidze et al. (2007) and Pontzer et al. (2010).

### 2.2. Phylogenetic analyses

Our sample comprises 105 cranial, mandibular, dental and postcranial characters collected from the above 16 terminal taxa (Table 1), with Dmanisi individuals treated as separate taxa to test their affinities.

Within species variation (polymorphism) was present in many characters for some species, but such characters have been argued to potentially contain phylogenetic information (Wiens 1995). Two approaches were used here to deal with intra-taxon variation. The first approach (dataset A) is the one most commonly used in phylogenetics analyses (CITE). Dataset A (SOM S1 and SOM Table S1) uses’multistate’ coding: taxa exhibiting polymorphism were coded as exhibiting all observed states which is the more typical approach in phylogenetics. The three situations previously discussed would be coded using a binary character, with taxa coded as states 0 (A), 0&1 (AB), and 1 (B). The second approach (dataset B) corresponds to the ‘scaled’ method in Wiens (1995), and makes the assumption that a lineage evolving from one fixed state to another should pass through a polymorphic intermediate stage. Dataset B uses’intermediate’ coding (SOM S2 and SOM Table S2): taxa exhibiting polymorphism were treated as exhibiting a distinct intermediate state between two alternative fixed states. Thus, taxa fixed for state A, taxa polymorphic for states A and B, and taxa fixed for state B were coded respectively as exhibiting states 0 (A), 1 (AB), and 2 (B) in an ordered 3-state character. This approach corresponds to the ‘scaled’ method in Wiens (1995), and makes the assumption that a lineage evolving from one fixed state to another should pass through a polymorphic intermediate stage.

Parsimony analyses were performed on datasets A (SOM S1 and B (SOM S2) using PAUP* (Swofford 2003). Character states are ordered where there is a clear linear morphocline (e.g., small, medium, large; see also discussion above regarding treatment of polymorphism).

Parsimony analyses placed the same weight for all character changes between fixed states (in unordered characters, or between adjacent states in ordered characters). Recoding of characters using intermediate coding (dataset B;) means changes between fixed states are indirectly weighted 2; to maintain parity of weighting, unordered multistate characters were upweighted to 2 in these analyses. Heuristic searches were employed with thorough settings: 1000 replicates, saving up to 1000 trees per replicate (to avoid memory overflow for searches which get trapped on very large islands of equally-parsimonious trees). All most-parsimonious trees were retained, and strict and majority-rule consensus trees constructed.

To infer apomorphies (derived character states) for clades of interest, characters were optimised using both accelerated transformation and delayed transformation.

Nonparametric bootstrapping (Felsenstein 1985; Hillis and Bull 1993), based on 200 replicates (with the heuristic search settings above), was used to evaluate topological support. The bootstrap frequencies of each clade on the most parsimonious tree were recorded, as well as the frequency in which all four Dmanisi individuals form a clade (see below).

We also evaluated the support for all Dmanisi taxa forming a single clade, consistent with the single-species hypothesis. In our main analyses, the Dmanisi individuals consistently separated into two groups: D4500_2600, which often appears near *Homo floresiensis*, and D2282, D2700_2735 and D2280, which often appear near *H. erectus, H. ergaster* and *H*.

*sapiens*. We thus tested the support for the alternative hypothesis that the Dmanisi taxa were a single taxon, by performing an analysis with all 4 Dmanisi individuals constrained to form a clade; this was done using datasets A and B. The nonparametric test of Templeton (1983) as implemented in PAUP* was also used to compare the scores of these alternative topologies.

We further examined the bootstrap frequency in which this alternative Dmanisi clade appeared in analyses when all relationships were unconstrained (all sampled clades and their frequencies are displayed in the default PAUP bootstrap output).

Finally, to complement the tree-based inferences, we looked at raw similarity between all taxa via the mean character difference (SOM Tables S3 and S4). This was performed using PAUP for both datasets A and B. In particular, we evaluated whether the Dmanisi individuals were all most phenetically similar to each other.

## 3. Results

As with many analyses dealing with morphology and fossils, topological support was often low compared to molecular/neontological thresholds (e.g. >70% bootstrap is often considered robust and roughly analogous to P<0.05); the results need to be viewed accordingly (see Discussion). The analysis using dataset A (multistate coding) retrieved a single shortest tree (271 steps, Fig. 1A), which groups three Dmanisi individuals together: D2280, D2282 and D2700_D2735, with bootstrap 43%; and D2280 and D2700_D2735 group together (bootstrap 48%). The Dmanisi clade is weakly placed as sister to *H. naledi, H. erectus*, *H. ergaster*, and *H. sapiens* (bootstrap 9%). However, the remaining Dmanisi individual (D4500_2600) grouped with *Homo floresiensis* (bootstrap 21%). A small but appreciable proportion of bootstrapped trees (7%) placed all 4 Dmanisi individuals together. The analysis where all four Dmanisi individuals were constrained to form a clade resulted in 4 shortest trees (274 steps; Fig 1B), which was not significantly different (Templeton test P>0.49 for all 4 trees) compared to the shortest tree (271 steps).

The analysis using dataset B (intermediate coding) retrieved two shortest trees (670 steps, Fig. 1C), where three Dmanisi individuals (D2280, D2282, D2700_D2735) again grouped together moderately strongly (bootstrap 69%); within this clade, D2280 and D2700_D2735 were again united (bootstrap 69%). This Dmanisi clade weakly clustered (bootstrap 15%) as sister to hominins *(H. habilis, H. erectus, H. ergaster*, and *H. sapiens*). Dmanisi individual D4500_2600 again grouped with *Homo floresiensis* (bootstrap 27%). A sizeable proportion of bootstrapped trees (16%) placed all 4 Dmanisi hominins together. The analysis in which all four Dmanisi individuals were constrained to form a clade resulted in one shortest tree, only two steps longer (Fig. 1D, 672 steps) than the unconstrained trees, and thus not significantly different (Templeton test P>0.76 compared to both best unconstrained trees). In this constrained tree, the clade of four Dmanisi individuals groups with *Homo floresiensis* (bootstrap 33%).

The character distances (SI 3) are also consistent with the above pattern. Three of the Dmanisi individuals (D2280, D2282, D2700_D2735) are highly similar (distances 0.3-0.44 in dataset A; 0.31-0.42 in dataset B). However, the remaining Dmanisi individual (D4500_2600) is much more divergent from the core three (distances 0.5-0.59 in dataset A and 0.5-0.57 in dataset B); in terms of character distances, D4500_2600 is actually more similar to many other hominins, including *H. floresiensis* (distance 0.41 dataset A; 0.42 dataset B).

## 4. Discussion

### 4.1 Phylogenetic results

When many taxa in a morphological phylogenetic analysis are very similar and differ only in subtle variation, statistically robust phylogenetic conclusions can be elusive. Furthermore, the extensive missing data in most fossils further reduces support. Both these factors are at work in the present study and the results should therefore be treated as heuristic: hypotheses that should be tested with additional data and new fossils.

Although the Dmanisi hominins have been variously assigned to *H. erectus* or *H. ergaster* (e.g. Gabunia and Vekua 1995; Rightmire et al. 2006; Pontzer et al. 2011), the Dmanisi hominins do not form sister taxa to *H. erectus (senso stricto)* or *H. ergaster* (included in *H. erectus* by some) in our analyses. There is thus no support in our dataset for the idea that any of the Dmanisi hominins are most closely related to (or conspecific) with *H. erectus or H. ergaster*.

Three of the four Dmanisi individuals form a clade (D2280, D2282, D2700_D2735), consistent with them being a single species. This clade is retrieved in all analyses, and the same three Dmanisi individuals are also highly similar according to character (phenetic) distances. Within this clade, D2280 and D2700_D2735 pair together strongly in all analyses.

However, it is less certain that Dmanisi individual D4500_2600 also belongs to this clade. In unconstrained phylogenetic analyses, the four Dmanisi individuals are separated into two clades/ D2280, D2280 and D2700_D2735 group with *H. habilis, H. erectus, H. ergaster*, and *H. sapiens*). The most compelling shared character supporting this relationship is a low symphyseal index (character 52: 1->2, consistency index = 1). D4500_D2600 groups with *H. floresiensis*, but there are no unique shared characters. There are also much larger character distances between D4500_2600 and the other three Dmanisi individuals (above, and SOM Table 3; SOM Table 4). Nevertheless, bootstrapping results (in unconstrained analyses) retrieve an appreciable frequency of trees that place all four Dmanisi individuals in a clade.

Furthermore, constrained analyses demonstrate that only three extra steps are required to’force’ all these taxa together. The most compelling character that support uniting D4500_2600 with the other three Dmanisi taxa is presence of anterior marginal tubercle (character 65, state change 0->1, Consistency Index=1; unique). However, it can be reasonably inferred that there is no evidence for more than two species in our analyses.

### 4.2 Implications of a single-species hypothesis for the Dmanisi hominins

Rightmire et al. (2017) interpret D4500_D2600 as a very robust male based upon its massive midface, prominent canine juga, and its high mandibular corpus with strong lateral and marginal relief. Its endocranial capacity is 546 +/-5 cm^3^ (Lordkipanidze et al. 2013), within the range of *Au. afarensis* (387cc-550cc; Schoenemann 2013), *Au. africanus* (435cc-560cc; Schoenemann 2013), and *Homo habilis* (509cc – 687cc; Schoenemann 2013). The D4500_D2600 endocranial capacity is 30% - 33% smaller than one of the other males in the group, D2280 (730 cm^3^), while a third adult male, D3444, has a brain size of 625 cm^3^ (Lordkipanidze et al. 2006). There is one assumed female in the assemblage: D2282 (Rightmire et al. 2017). D2282’s endocranial capacity is estimated to be 650-660 cm^3^ (Rightmire et al. 2006), 19%, larger than D4500. The subadult among the Dmanisi hominins has an endocranial capacity of between 600 cm^3^ and 612 cm^3^ (Rightmire et al. 2006), again, larger than D4500_D2600.

If we assume the Dmanisi hominins comprise a single species, we have an unexpected situation in which certain adult male crania may be notably smaller than other male crania and smaller than female crania. To our knowledge this pattern of dimorphism is not normal in other *Homo*, or in non-human primates. Rightmire et al. (2019) also observe that the pattern of putative sexual dimorphism among the Dmanisi population is surprising. They suggest that this pattern may differ from that in extant apes, living humans, and mid-Pleistocene hominins. The far-reaching implication of the patterns of sexual dimorphism observed in Dmanisi under a single species hypothesis for the hominins is that our definition of *Homo* must expand to include a pattern of sexual dimorphism in which some males in a species may have considerably smaller brains than females, and males, in that species. This pattern has not hitherto been reported in human evolution and would necessitate a considerable paradigm shift, and elucidation as to the evolutionary process behind such a pattern.

Also challenging for the single species hypothesis is how we may accommodate the range of variation in molar sequences among the Dmanisi hominins. In two mandibles M1 is larger than M2 and in one mandible the M1 is smaller than M2. The former characteristic is known primarily among Middle Pleistocene *Homo*, the latter occurs in Australopithecines, *Paranthropus*, and the majority of *H. habilis* and *H. rudolfensis.* Differences also extend to the M2/M3 sequences: M2 in D211 is larger than the M3, whereas D2600 shares the primitive pattern (M2 is smaller than M3), with the Australopiths, *Paranthropu*s and the majority of early *Homo* species. Under the single species model for the Dmanisi hominins we would need to accommodate a novel characteristic in *Homo*, in which Australopith/early *Homo* molar sequences and Middle-Pleistocene *Homo* molar sequences occur concurrently.

As for the mandibles, in D2735 and D211 the sagittal plane of the ramus forms an angle with the sagittal plane of the body and the two planes overlap (Fig. 2). Presumably, this mandibular architecture is the primitive state for the hominin clade. In D2600, however, the sagittal plane of the ramus and the sagittal plane of the body are practically parallel and hardly overlap —as, for example, it may be observed in Neanderthals. Presumably, this is a derived state for the hominin clade. To our knowledge, the primitive condition and the derived condition are not found simultaneously in the same hominin species.

**Fig. 2.**
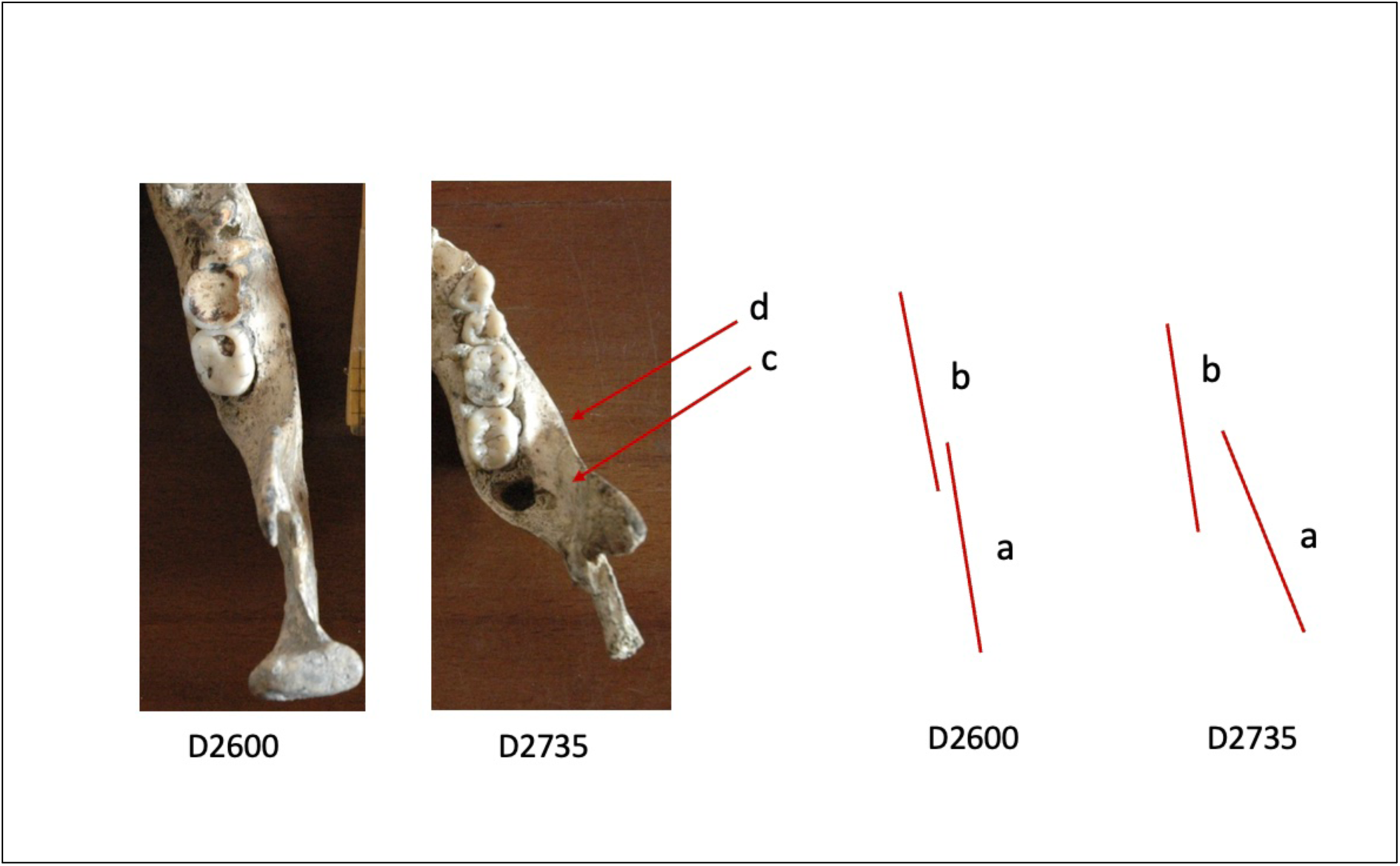
Dmanisi mandibles D2600 and D2735. (a) line contained in the sagittal plane of the mandibular ramus; (b) line contained in the sagittal plane of the mandibular body. See text for (c) and (d).

The lateral prominence (Figure 2, feature c) is advanced and more marked in D2735, although this specimen is smaller than D2600. Also, the width of the *sulcus extramolaris* (d) is greater in D2735 than in D2600. In D2735, the M3 is covered by the ramus when viewed in lateral view. This is not the case in D2600, where, in lateral view, the M3 is visible in its entirety.

We also note that two dietary regimes have been suggested to explain differences in wear patterns on D211’s and D 2735’s molars compared to the patterns identified on D2600.

Pontzer et al. (2011) suggest that the two individuals, D211 and D2735, were consuming fruits and tougher foods consistent with fracture-resistant foods such as meat or some underground storage organs (Pontzer et al. 2011) (e.g. roots and tubers). D2600, however, might have had a different dietary regime. Martín-Francés et al. (2014) propose that the extreme and rounded wear pattern observed in the anterior teeth of D2600 compared to its posterior teeth is similar to the pattern found on primarily folivorous species, such as chimpanzees and gorillas and would result from the intake of abrasive and fibrous foods such as plants and fruits. The dietary habits require pre-/para-masticatory preparation, such as gripping and stripping (Martín-Francés et al. 2014). No evidence for meat-eating or the consumption of underground food sources by D4500_D2600 is reported by Martín-Francés et al. (2014), while grasping and stripping cannot be inferred from observations on the occlusal wear on the incisors of D211 (Bermúdez de Castro, pers. comm.).

## Conclusions

The Dmanisi hominins are often included in *H. erectus* or *H. erectus* s. l. /*H. ergaster*, notwithstanding that they are formally designated as *Homo georgicus* with D4500_D2600 as the holotype. In our phylogenetic analyses, none of the Dmanisi hominins form a sister taxon to either *H. erectus* or *H. ergaster*. We hypothesise that the Dmanisi hominins did not share a unique common ancestor with *H. erectus* or *H. ergaster*, and we cannot support their attribution to either of those species.

Although all the Dmanisi hominins are attributed to a single species, the morphological variation evident among them has prompted questions about their heterogeneity with discussion focusing on whether one species or two are represented. We approached this question using phylogenetic analyses and by exploring further lines of evidence. Although our phylogenetic analyses did not lead us to propose two species among the Dmanisi hominins, there are nevertheless morphologically significant differences in the cranium, mandible and dentition of the individual represented by D4500_D2600 and the other Dmanisi hominins that are consistent with the view that D4500_D2600 represents a separate species. These are the unique and perplexing pattern of sexual dimorphism evident in the endocranial capacities of the assemblage when considered as one species; the dichotomy in mandibular molar size sequences; and in the presence of both a primitive and a derived form in the mandibular structures among the assemblage. We also note that D4500_D2600, in terms of character distances, is more similar to other hominins, including *H. floresiensis*, than it is to the other Dmanisi individuals.

We propose that the most parsimonious hypothesis for the Dmanisi hominins is that two species are present among the assemblage: *Homo georgicus* comprising D4500_D2600 and an un-named species comprising the other Dmanisi hominins: D2280, D2282_D211, D2700_D2735 and D3444_ D3900. The alternative hypothesis, that the assemblage comprises a single species, requires substantial paradigm shifts in our definition of *Homo*.

We surmise that the first hominin species at Dmanisi was *H. georgicus*, and that the species was probably present by 1.8 Ma. The other hominins, D2280, D2282_D211, D2700_D2735 and D3444_ D3900, accumulated at some time or times during the reverse polarity of 1.07 Ma and 1.77 Ma.

The specific ages of D2280, D2282_D211, D2700_D2735 and D3444_ D3900, however, remain unknown. Dating of the volcanic ashes in which each hominin was recovered, together with dating the ashes in the overlying strata to find the minimum date for the hominins, would likely produce a more refined understanding of the chronology for the Dmanisi hominins.

## Acknowledgements

We are most appreciative of previous research on the hominin remains from Dmanisi and the data that has been published. We thank Elen Feuerriegel for providing some postcranial data; Bradley Pillans, who advised on an interpretive matter related to the published geochronology at Dmanisi; and Philip Jones provided geological information to DA. In addition to co-authoring, José María Bermúdez de Castro supplied additional information about Dmanisi mandibular and dental features, and Maria Martinón-Torres contributed insights into the Dmanisi dentition and paleopathology. We appreciate the constructive comments provided by several peer reviewers. J.M. Bermúdez de Castro and M. Martinón-Torres are supported by the Project PID2021-122355NB-C33 financed by MCIN/ AEI/10.13039/501100011033/ FEDER, UE.

## Author contribution statement

DA initiated and coordinated the project. DA, JMBC, MMT provided the data, with some postcranial data being contributed by Dr Elen Feuerriegel. ML conducted the phylogenetic analyses, wrote the results and interpretation, and supplied the phylogenetic figures, tables and executable datasets. MBC and MMT took responsibility for the sections relating to mandibles and dentition. DA was responsible for writing the overall manuscript. All authors contributed revisions to the manuscript during the preparation process. All authors have provided approval for the manuscript to be submitted.

