## Supplementary S1 for "Where do the Dmanisi hominins fit on the human evolutionary tree?"

**S1. Character list with characters coded as multistate**

1. Parietal expansion. Relative biporionic width in relation to bi-parietal breadth; Biporionic breadth/ biparietal breadth x 100 (ordered).
    - 0. bi-parietal breadth very narrow in relation to biporionic breadth (index > 115)
    - 1. bi-parietal breadth relatively narrow in relation to biporionic (index; 103-115)
    - 2. bi-parietal somewhat wider or fairly similar in relation to biporionic breadth (index 84-100)
    - 3. bi-parietal breadth wider in relation to biporionic (index 70-78)
2. Maximum cranial breadth.
    - 0. at supramastoid region
    - 1. on parietals
3. Alveolar prognathism (ordered).
    - 0. very pronounced (< or = to 40 degrees)
    - 1. pronounced (pronounced 41-50 degrees)
    - 2. moderate projection (60 – 80 degrees)
    - 3. minimal or no projection (>80 degrees)
4. Inion relative to glabella (ordered).
    - 0. above
    - 1. on the same plane
    - 2. slightly below
    - 3. significantly below
5. Foramen spinosum (unordered).
    - 0. mostly absent
    - 1. on the sphenosquamosal suture
    - 2. contained within the greater wing of the sphenoid
6. Bregmatic eminence.
    - 0. absent.
    - 1. present
7. Basicranial flexion (ordered).
    - 0. not flexed or minimal flexion
    - 1. flexed (137–1380)
    - 2. strongly flexed (c. 1300)
8. Superior-inferior height of nuchal vs height of occipital (ordered).
    - 0. Superior-inferior height of nuchal greater than height of occipital
    - 1. nuchal and occipital of equal length
    - 2. Superior-inferior height of nuchal less than height of occipital
9. Posterior-anterior inclination of nuchal plane (ordered).
    - 0. steeply inclined
    - 1. weakly inclined
    - 2. horizontal
10. Muscle attachments on nuchal
    - 0. marked
    - 1. not marked
11. Supraorbital torus: flaring of trigones.
    - 0. present

- 1. absent

12. Occipital torus.
    - 0. present
    - 1. absent
13. Occipital sulcus.
    - 0. present
    - 1. absent
14. External occipital protuberance.
    - 0. prominent
    - 1. not prominent or absent
15. Mastoid processes inferior projection.
    - 0. does not project
    - 1. project below cranial base
16. Fissure between mastoid process and petrous crest of tympanic.
    - 0. present
    - 1. absent
17. Recess between the tympanic plate and entoglenoid pyramid.
    - 0. present
    - 1. absent
18. Vaginal process and crest (ordered).
    - 0. absent
    - 1. small
    - 2. extends along inferior surface of tube
    - 3. large
19. Medio-lateral angulation of articular eminence in basal view (ordered).
    - 0. angled medio-anteriorly
    - 1. parallel to coronal plane
    - 2. angled medio-posteriorly
20. Superior-inferior angulation of anterior wall of glenoid fossa (ordered).
    - 0. anterior wall horizontal
    - 1. oblique
    - 2. almost vertical
21. Robustness of tympanic (thickness of tympanic in norma lateralis, anterior edge of tympanic).
    - 0. thin (<2mm)
    - 1. thick (>2mm)
22. Postglenoid process (ordered).
    - 0. postglenoid process makes up much of the wall of the mandibular fossa.
    - 1. postglenoid and tympanic equally form the wall
    - 2. the tympanic makes up most of the wall
23. Mastoid process inflection.
    - 0. not inflected under cranial base
    - 1. inflected under cranial base
24. Mandibular/glenoid fossa overhang

Proportion of fossa that overhangs the external cranial vault.

- 0. >50%

- 1. <50%

25. Auditory meatus shape.
    - 0.  anteroposterorly compressed on vertical axis
    - 1. circular
26. Greatest axis of auditory meatus from inferior margin to superior margin (ordered).
    - 0. slopes anteriorly
    - 1. vertical
    - 2. slopes posteriorly
27. Tympanic trough.
    - 0. absent
    - 1. present
28. Supraorbital sulcus (ordered).
    - 0. present
    - 1. flat plane
    - 2. absent
29. Supraorbital torus.
    - 0. present
    - 1. absent
30. Zygomaticoalveolar crest (ordered).
    - 0. forms a full arch
    - 1. forms an arc
    - 2. relatively straight
31. Medial incursion of the superior temporal lines at coronal suture (ordered).
    - 0. crest or nearly so
    - 1. no inflection
    - 2. inflection
32. Metopic prominence.
    - 0. absent
    - 1. present
33. Form of glabella in superior view (ordered).
    - 0. depressed
    - 1. indistinct
    - 2. inflated
34. Location of infraorbital foramen.
    - 0. medially placed (closer to nasal aperture)
    - 1.  located mediolaterally under central lower orbit margin (or nearly so)
35. Canine juga.
    - 0. present
    - 1. absent
36. Anterior pillars.
    - 0. present
    - 1. absent
37. Condition of margo limitans.
    - 0. smooth curve
    - 1. forms a sill

38. Shape of clivus nas-oalveolaris medio-laterally at midsection of the naso-alveolar clivus (unordered).

- 0. bi-convex with central hollow
- 1. has a central spine
- 2. flat or slightly concave
- 3. convex and protrudes, or is rounded

39. Inferolateral nasal aperture margin.

- 0. blunt
- 1. sharp

40. Antero-inferior slope of malar in lateral view (ordered).

- 0. slopes anteriorly
- 1. vertical
- 2. slopes posteriorly

41. Flaring of zygomatic arch in superior view.

- 0. narrow divergence from lateral edge of cranium
- 1. diverges widely from lateral edge of cranium

42. Masseteric fossa (ordered).

- 0. flat
- 1. shallow
- 2. deep

43. Parietal keeling.

- 0. absent
- 1. present

44. Temporal line on the parietal (ordered).

- 0. fuses with nuchal line
- 1. continuity with supramastoid crest
- 2. no direct link with supramastoid crest

45. Obelionic depression.

- 0. absent
- 1. present

46. Angular torus.

- 0. absent
- 1. present

47. Divergence of tooth rows; lingual aspect (ordered).

- 0. converge posteriorly (and tooth rows are convex laterally)
- 1: parallel
- 2: diverge posteriorly

48. Palatine grooves and crests.

- 0. minimal or absent
- 1. moderate or marked

49. Depth of palate.

- 0. deep posteriorly
- 1. shallow posteriorly

50. Internal length of palate in relation to width (ordered).

- 0. very long (index 40-47)
- 1. long (index 53-60)

- 2. Intermediate (61-67)
- 3. Relatively short in relation to length (>70)

51. Symphyseal region (unordered).
    - 0. no mental protuberance and retreats
    - 1. no mental protuberance and vertical
    - 2. slight symphyseal ridge or swelling and retreats
    - 3. mental protuberance with keel (chin) present
52. Symphysis anterior-posterior width in relation to external arch breadth at M2 (ordered)
    - 0. Index: >35
    - 1. Index: 30-35.0
    - 2. Index: 24.2-30.0
53. Ramus obscures M3 .
    - 0. yes, at least partly
    - 1. no
54. Relationship between corpus and ramus.
    - 0. Minimal or weak acute angle on alveolar border at M3
    - 1. Internal alveolar border turns strongly buccalwise at M3 forming a strong acute angle
55. Position of mental foramina (ordered)
    - 0. under or anterior to P3
    - 1. under septum P3 P4
    - 2. under P4
    - 3. under P4 M1 septum
    - 4. under M1
56. Number of mental foramina.
    - 0. single
    - 1. multiple
57. Deepest part of sigmoid notch (ordered)
    - 0. towards coronoid
    - 1. centrally
    - 2. towards condyle
58. Marginal torus.
    - 0. absent
    - 1. present
59. Position of lateral prominence (ordered)
    - 0. below M1
    - 1. below M2
    - 2. between M2/M3
60. Mid-ramus ‘waisting’.
    - 0. absent
    - 1. present
61. Superior lateral torus.
    - 0. absent
    - 1. narrow
62. Superior transverse torus.
    - 0. prominent to weak

- 1. very weak or absent

63. Inferior transverse torus (ordered).
    - 0. prominent
    - 1. weak
    - 2. absent
64. Submandibular fossa
    - 0. shallow
    - 1. deep
65. Anterior marginal tubercle
    - 0. absent
    - 1. present
66. P3 root morphology (ordered).
    - 0. two roots and diverge near crown
    - 1. two roots and diverge at middle of root
    - 2. Tomes root
    - 3. single root
67. M1 fissure pattern.
    - 0. Y contact between metaconid and hypoconid
    - 1. + point contact between metaconid, hypoconid, protoconid, entoconid
68. M2 fissure pattern
    - 0. Y contact between metaconid and hypoconid
    - 1. + point contact between metaconid, hypoconid, protoconid, entoconid
69. M3 fissure pattern.
    - 0. Y contact between metaconid and hypoconid
    - 1. + point contact between metaconid, hypoconid, protoconid, entoconid
70. Molars: M3 tuberculum sextum.
    - 0. absent
    - 1. present
71. M3, C7 (ordered).
    - 0. C7 absent
    - 1. a short fissure is present branching off the lingual fissure into the metaconid, but no cusp can be identified
    - 2. a longer and more definite fissure runs off the lingual fissure, with a small cusp-like formation identifiable between them
    - 3. fissures clearly demarcate a well-developed accessory cusp
72. P4 cusp number (ordered)
    - 0. two cusps
    - 1. three cusps
    - 2. four cusps
73. Prominence of lingual ridge of lower canine.
    - 0. prominent
    - 1. not prominent
74. P4 metaconid development (ordered).
    - 0. strong
    - 1. weak
    - 2. absent

75. P3 talonid height in relation to protoconid (ordered).
    - 0. very low
    - 1. low
    - 2. moderate
76. P3 occlusal crown outline.
    - 0. along axis of tooth row
    - 1. at angle to tooth row
77. P3 metaconid.
    - 0. present
    - 1. absent
78. Mid-trigonid crest M1.
    - 0. present
    - 1. rare or absent
79. M3;M2 size ratios.
    - 0. M3> M2
    - 1. M3</= M2
80. Infraglenoid tubercle, scapula.
    - 0. broad
    - 1. narrow
81. Axillary border (unordered).
    - 0. pronounced and rounded
    - 1. flattened
    - 2. sharp ridge
82. Spinous process root, scapula.
    - 0. robust
    - 1. gracile
83. Orientation of scapula spine (ordered).
    - 0. 20º–40º
    - 1. 41º–60º
    - 2. >60º
84. Bar-glenoid angle.
    - 0. <135 º (superiorly directed glenoid fossa)
    - 1. >150 º
85. Infra spinous fossa, scapula.
    - 0. Flat or mild concavity with abrupt step to axillary border
    - 1. Convex with gentle curve to axillary border
86. Anteroposterior shaft curvature, femur.
    - 0. retroflexed
    - 1. straight
87. Humeral torsion.
    - 0. high
    - 1. low
88. Linea aspera.
    - 0. not well-developed
    - 1. well-developed
89. Intertrochanteric crest.

- 0. flattened
- 1. strongly developed

90. Greater trochanter.
    - 0. flat lateral apophysis
    - 1. prominent lateral apophysis
91. Biomechanical neck length, cf femoral head diameter.
    - 0. short
    - 1. long
92. Femur shaft in cross-section (unordered).
    - 0. circular
    - 1. oval
    - 2. tear-shaped
93. Pilaster on femur shaft (ordered).
    - 0. absent
    - 1. present but weakly defined
    - 2. well-defined
94. Bicondylar angle
    - 0. low
    - 1. high
95. Robustness of fibula (ordered).
    - 0. high
    - 1. medium
    - 2. low
96. Robustness of tibia
    - 0. high
    - 1. low
97. Ilium.
    - 0. Moderate flaring
    - 1. flares strongly beyond margin of acetabulum
98. Iliac pillar (ordered)
    - 0. virtually no pillar
    - 1. pillar anterior and weak
    - 2. pillar more posterior and marked
99. Postcranial proportions (ordered)
    - 0. humerofemoral index high (>80)
    - 1. humerofemoral index medium (61–80)
    - 2. humerofemoral index low (<60)
100. Tibia torsion (ordered).
    - 0. negative torsion
    - 1. near zero
    - 2. positive torsion
101. Tarsus: talar head torsion
    - 0. low torsion
    - 1. high torsion
102. Navicular tuberosity  (ordered).
    - 0. large (tuberosities project beyond facet for entocuneiform)

- 1. medium projection of tuberosities
- 2. reduced (tuberosities do not reach facet for entocuneiform)

103. Femoral neck index (ratio of AP/SI)

- 0. high index
- 1. low index

104. Relative clavicle length (ordered).

- 0. short
- 1. medium
- 2. long
- 4. long

105. Cranial capacity (ordered).

- 0. small (<560cc)
- 1. medium (>560cc – 700cc)
- 2. large (>700cc)
