## Supplementary S2 for "Where do the Dmanisi hominins fit on the human evolutionary tree?"

**S2. Character list with characters coded as intermediate**

1. Parietal expansion. Relative biporionic width in relation to bi-parietal breadth; Biporionic breadth/ biparietal breadth x 100 (ordered).
- 0. bi-parietal breadth very narrow in relation to biporionic breadth (index > 115)
- 1. states 0 & 2
- 2. bi-parietal breadth relatively narrow in relation to biporionic (index; 103-115)
- 3. states 2 & 4
- 4. bi-parietal somewhat wider or fairly similar in relation to biporionic breadth (index 84-100)
- 5. states 4 & 6
- 6. bi-parietal breadth wider in relation to biporionic (index 70-78)

2. Maximum cranial breadth (ordered).
- 0. at supramastoid region
- 1. states 0 & 2
- 2. on parietals

3. Alveolar prognathism (ordered).
- 0. very pronounced (< or = to 40 degrees)
- 1. states 0 & 2
- 2. pronounced (pronounced 41-50 degrees)
- 3. states 2 & 4
- 4. moderate projection (60 – 80 degrees)
- 5. states 4 & 6
- 6. minimal or no projection (>80 degrees)

4. Inion relative to glabella (ordered).
- 0. above
- 1. states 0 & 2
- 2. on the same plane
- 3. states 2 & 4
- 4. slightly below
- 5. states 4 & 6
- 6. significantly below

5. Foramen spinosum (unordered).
- 0. mostly absent
- 1. on the sphenosquamosal suture
- 2. contained within the greater wing of the sphenoid

6. Bregmatic eminence (ordered).
- 0. absent.
- 1. states 0 & 2
- 2. present

7. Basicranial flexion (ordered).
- 0. not flexed or minimal flexion
- 1. states 0 & 2
- 2. flexed (137–1380)
- 3. states 2 & 4
- 4. strongly flexed (c. 1300)

8. Superior-inferior height of nuchal vs height of occipital (ordered).
- 0. Superior-inferior height of nuchal greater than height of occipital
- 1. states 0 & 2
- 2. nuchal and occipital of equal length
- 3. states 2 & 4
- 4. Superior-inferior height of nuchal less than height of occipital

9. Posterior-anterior inclination of nuchal plane (ordered).
- 0. steeply inclined
- 1. states 0 & 2
- 2. weakly inclined
- 3. states 2 & 4
- 4. horizontal

10. Muscle attachments on nuchal (ordered).
- 0. marked
- 1. states 0 & 2
- 2. not marked

11. Supraorbital torus: flaring of trigones (ordered).
- 0. present
- 1. states 0 & 2
- 2. absent

12. Occipital torus (ordered).
- 0. present
- 1. states 0 & 2
- 2. absent

13. Occipital sulcus (ordered).
- 0. present
- 1. states 0 & 2
- 2. absent

14. External occipital protuberance (ordered).
- 0. prominent
- 1. states 0 & 2
- 2. not prominent or absent

15. Mastoid processes inferior projection (ordered).
- 0. does not project
- 1. states 0 & 2
- 2. project below cranial base

16. Fissure between mastoid process and petrous crest of tympanic (ordered).
- 0. present
- 1. states 0 & 2
- 2. absent

17. Recess between the tympanic plate and entoglenoid pyramid (ordered).
- 0. present
- 1. states 0 & 2
- 2. absent

18. Vaginal process and crest (ordered).
- 0. absent

- 1. states 0 & 2
- 2. small
- 3. states 2 & 4
- 4. extends along inferior surface of tube
- 5. states 4 & 6
- 6. large

19. Medio-lateral angulation of articular eminence in basal view (ordered).
    - 0. angled medio-anteriorly
    - 1. states 0 & 2
    - 2. parallel to coronal plane
    - 3. states 2 & 4
    - 4. angled medio-posteriorly
20. Superior-inferior angulation of anterior wall of glenoid fossa (ordered).
    - 0. anterior wall horizontal
    - 1. states 0 & 2
    - 2. oblique
    - 3. states 2 & 4
    - 4. almost vertical
21. Robustness of tympanic, thickness of tympanic in norma lateralis, anterior edge of tympanic (ordered).
    - 0. thin (<2mm)
    - 1. states 0 & 2
    - 2. thick (>2mm)
22. Postglenoid process (ordered).
    - 0. postglenoid process makes up much of the wall of the mandibular fossa.
    - 1. states 0 & 2
    - 2. postglenoid and tympanic equally form the wall
    - 3. states 2 & 4
    - 4. the tympanic makes up most of the wall
23. Mastoid process inflection (ordered).
    - 0. not inflected under cranial base
    - 1. states 0 & 2
    - 2. inflected under cranial base
24. Mandibular/glenoid fossa overhang, proportion of fossa that overhangs the external cranial vault (ordered).
    - 0. >50%
    - 1. states 0 & 2
    - 2. <50%
25. Auditory meatus shape (ordered).
    - 0. anteroposterorly compressed on vertical axis
    - 1. states 0 & 2
    - 2. circular
26. Greatest axis of auditory meatus from inferior margin to superior margin (ordered).
    - 0. slopes anteriorly
    - 1. states 0 & 2
    - 2. vertical

- 3. states 2 & 4
- 4. slopes posteriorly

27. Tympanic trough (ordered).
    - 0. absent
    - 1. states 0 & 2
    - 2. present
28. Supraorbital sulcus (ordered).
    - 0. present
    - 1. states 0 & 2
    - 2. flat plane
    - 3. states 2 & 4
    - 4. absent
29. Supraorbital torus (ordered).
    - 0. present
    - 1. states 0 & 2
    - 2. absent
30. Zygomaticoalveolar crest (ordered).
    - 0. forms a full arch
    - 1. states 0 & 2
    - 2. forms an arc
    - 3. states 2 & 4
    - 4. relatively straight
31. Medial incursion of the superior temporal lines at coronal suture (ordered).
    - 0. crest or nearly so
    - 1. states 0 & 2
    - 2. no inflection
    - 3. states 2 & 4
    - 4. inflection
32. Metopic prominence (ordered).
    - 0. absent
    - 1. states 0 & 2
    - 2. present
33. Form of glabella in superior view (ordered).
    - 0. depressed
    - 1. states 0 & 2
    - 2. indistinct
    - 3. states 2 & 4
    - 4. inflated
34. Location of infraorbital foramen (ordered).
    - 0. medially placed (closer to nasal aperture)
    - 1. states 0 & 2
    - 2. located mediolaterally under central lower orbit margin (or nearly so)
35. Canine juga (ordered).
    - 0. present
    - 1. states 0 & 2
    - 2. absent

36. Anterior pillars (ordered).
- 0. present
- 1. states 0 & 2
- 2. absent

37. Condition of margo limitans (ordered).
- 0. smooth curve
- 1. states 0 & 2
- 2. forms a sill

38. Shape of clivus nas-oalveolaris medio-laterally at midsection of the naso-alveolar clivus (unordered).
- 0. bi-convex with central hollow
- 1. has a central spine
- 2. flat or slightly concave
- 3. convex and protrudes, or is rounded

39. Inferolateral nasal aperture margin (ordered).
- 0. blunt
- 1. states 0 & 2
- 2. sharp

40. Antero-inferior slope of malar in lateral view (ordered).
- 0. slopes anteriorly
- 1. states 0 & 2
- 2. vertical
- 3. states 2 & 4
- 4. slopes posteriorly

41. Flaring of zygomatic arch in superior view (ordered).
- 0. narrow divergence from lateral edge of cranium
- 1. states 0 & 2
- 2. diverges widely from lateral edge of cranium

42. Masseteric fossa (ordered).
- 0. flat
- 1. states 0 & 2
- 2. shallow
- 3. states 2 & 4
- 4. deep

43. Parietal keeling (ordered).
- 0. absent
- 1. states 0 & 2
- 2. present

44. Temporal line on the parietal (ordered).
- 0. fuses with nuchal line
- 1. states 0 & 2
- 2. continuity with supramastoid crest
- 3. states 2 & 4
- 4. no direct link with supramastoid crest

45. Obelionic depression (ordered).
- 0. absent

- 1. states 0 & 2
- 2. present

46. Angular torus (ordered).
    - 0. absent
    - 1. states 0 & 2
    - 2. present
47. Divergence of tooth rows; lingual aspect (ordered).
    - 0. converge posteriorly (and tooth rows are convex laterally)
    - 1. states 0 & 2
    - 2: parallel
    - 3. states 2 & 4
    - 4: diverge posteriorly
48. Palatine grooves and crests (ordered).
    - 0. minimal or absent
    - 1. states 0 & 2
    - 2. moderate or marked
49. Depth of palate (ordered).
    - 0. deep posteriorly
    - 1. states 0 & 2
    - 2. shallow posteriorly
50. Internal length of palate in relation to width (ordered).
    - 0. very long (index 40-47)
    - 1. states 0 & 2
    - 2. long (index 53-60)
    - 3. states 2 & 4
    - 4. Intermediate (61-67)
    - 5. states 4 & 6
    - 6. Relatively short in relation to length (>70)
51. Symphyseal region (unordered).
    - 0. no mental protuberance and retreats
    - 1. no mental protuberance and vertical
    - 2. slight symphyseal ridge or swelling and retreats
    - 3. mental protuberance with keel (chin) present
52. Symphysis anterior-posterior width in relation to external arch breadth at M2 (ordered)
    - 0. Index: >35
    - 1. states 0 & 2
    - 2. Index: 30-35.0
    - 3. states 2 & 4
    - 4. Index: 24.2-30.0
53. Ramus obscures M3 (ordered).
    - 0. yes, at least partly
    - 1. states 0 & 2
    - 2. no
54. Relationship between corpus and ramus (ordered).
    - 0. Minimal or weak acute angle on alveolar border at M3
    - 1. states 0 & 2

- 2. Internal alveolar border turns strongly buccalwise at M3 forming a strong acute angle

55. Position of mental foramina (ordered)
    - 0. under or anterior to P3
    - 1. states 0 & 2
    - 2. under septum P3 P4
    - 3. states 2 & 4
    - 4. under P4
    - 5. states 4 & 6
    - 6. under P4 M1 septum
    - 7. states 6 & 8
    - 8. under M1
56. Number of mental foramina (ordered).
    - 0. single
    - 1. states 0 & 2
    - 2. multiple
57. Deepest part of sigmoid notch (ordered)
    - 0. towards coronoid
    - 1. states 0 & 2
    - 2. centrally
    - 3. states 2 & 4
    - 4. towards condyle
58. Marginal torus (ordered).
    - 0. absent
    - 1. states 0 & 2
    - 2. present
59. Position of lateral prominence (ordered)
    - 0. below M1
    - 1. states 0 & 2
    - 2. below M2
    - 3. states 2 & 4
    - 4. between M2/M3
60. Mid-ramus ‘waisting’ (ordered).
    - 0. absent
    - 1. states 0 & 2
    - 2. present
61. Superior lateral torus (ordered).
    - 0. absent
    - 1. states 0 & 2
    - 2. narrow
62. Superior transverse torus (ordered).
    - 0. prominent to weak
    - 1. states 0 & 2
    - 2. very weak or absent
63. Inferior transverse torus (ordered).
    - 0. prominent

- 1. states 0 & 2
- 2. weak
- 3. states 2 & 4
- 4. absent

64. Submandibular fossa (ordered).
    - 0. shallow
    - 1. states 0 & 2
    - 2. deep
65. Anterior marginal tubercle (ordered).
    - 0. absent
    - 1. states 0 & 2
    - 2. present
66. P3 root morphology (ordered).
    - 0. two roots and diverge near crown
    - 1. states 0 & 2
    - 2. two roots and diverge at middle of root
    - 3. states 2 & 4
    - 4. Tomes root
    - 5. states 4 & 6
    - 6. single root
67. M1 fissure pattern (ordered).
    - 0. Y contact between metaconid and hypoconid
    - 1. states 0 & 2
    - 2. + point contact between metaconid, hypoconid, protoconid, entoconid
68. M2 fissure pattern (ordered).
    - 0. Y contact between metaconid and hypoconid
    - 1. states 0 & 2
    - 2. + point contact between metaconid, hypoconid, protoconid, entoconid
69. M3 fissure pattern (ordered).
    - 0. Y contact between metaconid and hypoconid
    - 1. states 0 & 2
    - 2. + point contact between metaconid, hypoconid, protoconid, entoconid
70. Molars: M3 tuberculum sextum (ordered).
    - 0. absent
    - 1. states 0 & 2
    - 2. present
71. M3, C7 (ordered).
    - 0. C7 absent
    - 1. states 0 & 2
    - 2. a short fissure is present branching off the lingual fissure into the metaconid, but no cusp can be identified
    - 3. states 2 & 4
    - 4. a longer and more definite fissure runs off the lingual fissure, with a small cusp-like formation identifiable between them
    - 5. states 4 & 6
    - 6. fissures clearly demarcate a well-developed accessory cusp

72. P4 cusp number (ordered)
    - 0. two cusps
    - 1. states 0 & 2
    - 2. three cusps
    - 3. states 2 & 4
    - 4. four cusps
73. Prominence of lingual ridge of lower canine (ordered).
    - 0. prominent
    - 1. states 0 & 2
    - 2. not prominent
74. P4 metaconid development (ordered).
    - 0. strong
    - 1. states 0 & 2
    - 2. weak
    - 3. states 2 & 4
    - 4. absent
75. P3 talonid height in relation to protoconid (ordered).
    - 0. very low
    - 1. states 0 & 2
    - 2. low
    - 3. states 2 & 4
    - 4. moderate
76. P3 occlusal crown outline (ordered).
    - 0. along axis of tooth row
    - 1. states 0 & 2
    - 2. at angle to tooth row
77. P3 metaconid (ordered).
    - 0. present
    - 1. states 0 & 2
    - 2. absent
78. Mid-trigonid crest M1 (ordered).
    - 0. present
    - 1. states 0 & 2
    - 2. rare or absent
79. M3;M2 size ratios (ordered).
    - 0. M3> M2
    - 1. states 0 & 2
    - 2. M3</= M2
80. Infraglenoid tubercle, scapula (ordered).
    - 0. broad
    - 1. states 0 & 2
    - 2. narrow
81. Axillary border (unordered).
    - 0. pronounced and rounded
    - 1. flattened
    - 2. sharp ridge

82. Spinous process root, scapula (ordered).
    - 0. robust
    - 1. states 0 & 2
    - 2. gracile
83. Orientation of scapula spine (ordered).
    - 0. 20º–40º
    - 1. states 0 & 2
    - 2. 41º–60º
    - 3. states 2 & 4
    - 4. >60º
84. Bar-glenoid angle.
    - 0. <135 º (superiorly directed glenoid fossa)
    - 1. states 0 & 2
    - 2. >150 º
85. Infra spinous fossa, scapula (ordered).
    - 0. Flat or mild concavity with abrupt step to axillary border
    - 1. states 0 & 2
    - 2. Convex with gentle curve to axillary border
86. Anteroposterior shaft curvature, femur (ordered).
    - 0. retroflexed
    - 1. states 0 & 2
    - 2. straight
87. Humeral torsion (ordered).
    - 0. high
    - 1. states 0 & 2
    - 2. low
88. Linea aspera (ordered).
    - 0. not well-developed
    - 1. states 0 & 2
    - 2. well-developed
89. Intertrochanteric crest (ordered).
    - 0. flattened
    - 1. states 0 & 2
    - 2. strongly developed
90. Greater trochanter (ordered).
    - 0. flat lateral apophysis
    - 1. states 0 & 2
    - 2. prominent lateral apophysis
91. Biomechanical neck length, cf femoral head diameter (ordered).
    - 0. short
    - 1. states 0 & 2
    - 2. long
92. Femur shaft in cross-section (unordered).
    - 0. circular
    - 1. oval
    - 2. tear-shaped

93. Pilaster on femur shaft (ordered).
- 0. absent
- 1. states 0 & 2
- 2. present but weakly defined
- 3. states 2 & 4
- 4. well-defined

94. Bicondylar angle (ordered).
- 0. low
- 1. states 0 & 2
- 2. high

95. Robustness of fibula (ordered).
- 0. high
- 1. states 0 & 2
- 2. medium
- 3. states 2 & 4
- 4. low

96. Robustness of tibia (ordered).
- 0. high
- 1. states 0 & 2
- 2. low

97. Ilium (ordered).
- 0. Moderate flaring
- 1. states 0 & 2
- 2. flares strongly beyond margin of acetabulum

98. Iliac pillar (ordered)
- 0. virtually no pillar
- 1. states 0 & 2
- 2. pillar anterior and weak
- 3. states 2 & 4
- 4. pillar more posterior and marked

99. Postcranial proportions (ordered)
- 0. humerofemoral index high (>80)
- 1. humerofemoral index medium (61–80)
- 2. humerofemoral index low (<60)

100. Tibia torsion (ordered).
- 0. negative torsion
- 1. states 0 & 2
- 2. near zero
- 3. states 2 & 4
- 4. positive torsion

101. Tarsus: talar head torsion (ordered).
- 0. low torsion
- 1. states 0 & 2
- 2. high torsion

102. Navicular tuberosity (ordered).
- 0. large (tuberosities project beyond facet for entocuneiform)

- 1. states 0 & 2
- 2. medium projection of tuberosities
- 3. states 2 & 4
- 4. reduced (tuberosities do not reach facet for entocuneiform)

103. Femoral neck index, ratio of AP/SI (ordered).

- 0. high index
- 1. states 0 & 2
- 2. low index

104. Relative clavicle length (ordered).

- 0. short
- 1. states 0 & 2
- 2. medium
- 3. states 2 & 4
- 4. long

105. Cranial capacity (ordered).

- 0. small (<560cc)
- 1. states 0 & 2
- 2. medium (>560cc – 700cc)
- 3. states 2 & 4.
- 4. large (>700cc)
